# Influenza-induced alveolar macrophages protect against death by malaria-associated acute lung injury

**DOI:** 10.1101/2024.07.12.601219

**Authors:** Jenna S. Reed, Ritika Nayan, Margot Deckers, Brian D. Evavold, Tracey J. Lamb

**Affiliations:** Department of Pathology, University of Utah, 15 N Medical Drive, Salt Lake City 84112, US; KU Leuven Department of Microbiology, Immunology and Transplantation, Rega - Herestraat 49 - box 1044, 3000 Leuven, Belgium

**Keywords:** Malaria, malaria-associated acute lung injury, influenza, co-infection, arginase, myeloid cells

## Abstract

Lower respiratory tract infections are common in malaria-endemic areas, and there is some evidence that co-infections between various bacteria/viruses and *Plasmodium* may affect disease prognosis. In this study, we report the novel finding that co-infection with influenza/A/X31 protects mice from death by *Plasmodium berghei* NK65-Edinburgh, a model of severe malarial pulmonary leak which underpins malaria-associated acute lung injury (MA-ALI) and malaria-associated acute respiratory distress (MA-ARDS). Co-infected mice exhibit equivalent parasitemia as mice with malaria only, suggesting that the survival phenotype is due to differences in immune kinetics. We demonstrated that the pulmonary leak typical of *Pb*E is attenuated in co-infected mice without alteration in CD8 T cell activation and recruitment to the lung. Upon further examination of the immune response to influenza/A/X31 we identified a population of arginase 1-expressing alveolar macrophages that traffic to the lungs early during infection. In vitro these macrophages inhibit CD8 T cell activation and proliferation better than non-arginase expressing cells. Removal of arginase-1 expressing alveolar macrophages *in vivo* via administration of the antimetabolite gemcitabine removed the protective effects of influenza/A/X31co-infection on MA-ALI. This study opens a route to better understanding of how to modulate the immunopathology observed in pulmonary leak associated with severe malaria, which must be achieved to rationally design therapeutic interventions for MA-ARDS / MA-ALI.

## Introduction

Lower respiratory tract infections are common in malaria endemic areas. In 2019, there were a reported 489 million lower respiratory infections, making them the 4^th^ leading cause of death amongst all ages and the leading cause in children under 5 years old.^1^ These severe cases disproportionately occur in malaria endemic areas, but the causes underpinning this observation are not currently understood.^2^ It has been suggested that *Plasmodium*-infected individuals may have elevated susceptibility to invasive bacterial and viral diseases.^3–9^ Viral coinfections such as influenza are prevalent but under-reported and understudied in malaria-endemic regions. The effects of malaria-influenza coinfection are currently unknown.

Malaria-associated acute respiratory distress syndrome (MA-ARDS) and malaria-associated acute lung injury (MA-ALI) are severe manifestations of malaria that results in pulmonary vascular leak.^2,10–12^ MA-ARDS/MA-ALI can result from all five human-infecting species of *Plasmodium*,^1^ but is a common feature of pathology in adults infected with *P. vivax*^13^ or *P. knowlesi*.^2,10^ MA-ARDS/MA-ALI is associated with alveolar inflammation in response to sequestered *Plasmodium*-infected red blood cells (iRBCs) leading to breakdown in the integrity of the alveolar capillary membranes and vascular leakage of plasma fluid into the interstitium and alveoli.^11,12,14,15^ There is also accumulation of leukocytes in the microvessels.^16^ Mouse models of malaria-associated organ pathologies demonstrate the requirement for CD8 T cells in mediating vascular leak at the pulmonary and blood-brain barriers.^14,15^ The exact mechanisms by which CD8 T cells cause vascular leak in C57BL/6 mice infected with *Plasmodium berghei* NK65-Edinburgh (*Pb*E) are poorly understood but likely involve a combination of cytolysis of pulmonary endothelial cells^1^ and down regulation of junction proteins,^17,18^ disrupting the integrity of the pulmonary vasculature.

Despite the severe manifestations of MA-ARDS/MA-ALI, the only clinical treatment available is positive ventilation therapy in combination with anti-malarial chemotherapies. These treatments are ineffective, and there is still an unacceptably high mortality rate of 50% after treatment.^10,15^ As such, it is crucial to find novel strategies that will work in conjunction with anti-parasitic drugs to disrupt pulmonary pathology. Here we show that a non-lethal influenza A/X31 co-infection protects against pathology in a mouse model of MA-ALI / MA-ARDS. Protection was dependent on the timing of co-infection and influenza/A/X31 was only protective when occurring in the early part of *Pb*E infection, suggesting an innate immune system-mediated protective effect. Influenza/A/X31 induces an influx of arginase 1-expressing monocytes to the lung and was needed for protection against MA-ALI / MA-ARDS via suppressive effects on *Pb*E-reactive CD8 T cells. This data represents a potential new strategy for therapeutic targeting for MA-ALI / MA-ARDS.

## Material and methods

### Mice and infections

All animal work was performed with permission from the University of Utah Institutional Animal Care and Use Committee (Protocol #00002078). Female C57BL/6J mice aged 6-8 weeks (Line #000664, Jackson Laboratories) and were infected by intraperitoneal (i.p.) injection of 5×10^5^ iRBCs *Plasmodium berghei* NK65-Edinburgh (*Pb*E) (a kind gift of Dr. Philippe van den Steen, KU Leuven, Belgium). Influenza/A/X31 (a kind gift from Dr. Jacob Kohlmeier, Emory University, USA) was administered intranasally (i.n.) under isoflurane anesthesia with 3×10^4^ plaque forming units (PFU) of influenza/A/X31 per mouse.

We used a published clinical scoring system to monitor mice for MA-ARDS^19^. Briefly each mouse was monitored for social activity deficits (SA, 0=absent, 1=mild, 2=severe), limb grasp (LG, 0=normal, 1=mild deficit, 2=severe deficit), body tone (BT, 0=normal, 1=mild deficit, 2=severe deficit), piloerection (PE, 0=absent, 1=present), shivering (Sh, 0=absent, 1=present), abnormal breathing (AB, 0=absent, 1=present), dehydration (D, 0=absent, 1=mild, 2=severe), incontinence (I, 0=absent, 1=mild, 2=severe), and paralysis (P, 0=absent, 1=mild, 2=severe). The final clinical score was calculated as SA+LG+BT+TC+PE+Sh+AB+(3*(D+I+P)). Mice were weighed on a top pan balance and anemia measured on a Beckman ZI Coulter Counter. Parasitemia was evaluated using flow cytometry with the addition of CD45-APC (1:200, Biolegend #109813), CD71-PE (1:200, Biolegend #113807), Hoechst 34580 (1:200, BD Biosciences #565877). Samples were analyzed on a BD Biosciences X20 analyzer with collection of 100,000 events. RBCs were identified as CD45^-^, while Hoechst^+^ cells were considered pRBCs.

### Vascular permeability assays

Mice were injected intraperitoneally with 100 µL of 1% solution of Evans blue dye (Sigma #314-13-6) or 2 mg of 70 kDA fluorescein (FITC)-conjugated dextran (Thermofisher #D1823) in sterile PBS at day 6 post infection with *P. berghei* NK65e and euthanized for analysis after 1 hour. To assess leak by Evan’s blue the lungs were harvested and imaged with a USB desktop digital microscope (Veho) before being placed in 1 mL N,N-dimethylformamide (Sigma #68-12-2) an incubated for 4 days at 37°C to extract the dye from the tissue. The dye concentration in the supernatant was then quantified on a plate reader (Biotek Synergy) at an absorbance of 620 nm and quantified by comparing to a standard curve. To assess leak by FITC-Dextran bronchoalveolar lavage fluid (BALF) was collected from mice by inserting an 18 G catheter into the trachea and flushing the lungs twice with 750 µL 1X PBS each time (1.5 mL total) and fluorescence (Ex/Em 482/525) quantified on a plate reader (Biotek Synergy).

### MicroBCA Assay

Following euthanasia at 6 dpi, BALF was centrifuged at 2000 x g for 10 minutes and protein concentration of BALF quantified using a micro Bicinchoninic Acid (BCA) assay (Thermofisher Scientific #23235). The color change indicating protein concentration was quantified using a plate reader (Biotek Synergy) at an absorbance of 562 nm relative to a bovine serum albumin (BSA) standard curve.

### Isolation of single cell suspensions

Tissues were collected in RPMI (iRPMI; RPMI 1640 supplemented with 200 mM L-glutamine, 1X penicillin/streptomycin + 4 mM L-glutamine). Single cell suspensions from lung tissue obtained using a Miltenyi gentleMACS machine in 5 mL digestion media (5 mL iRPMI, 2.5 mg Collagenase D, 2000 U DNase I) and filtered through a 100 μm cell strainer. Spleens were homogenized by aseptically pushing the tissue through a 100 μm cell strainer. Homogenates were centrifuged at 300 x g for 5 minutes and red blood cells removed by incubating with 1X RBC Lysis Buffer (eBioscience 00-4333-57) for 1 minute at room temperature before dilution with 9 mL cold complete RPMI (cRPMI; RPMI 1640 supplemented with 200 mM L-glutamine (Corning 10-040-CV, 10% FBS, 1X penicillin/streptomycin + 4 mM L-glutamine, 0.01 M HEPES, 0.05 mM 2-β-mercaptoethanol). Following centrifugation and removal of the supernatant, homogenates were resuspended in cRPMI and filtered again.

### Flow cytometry

2×10^6^ cells per sample were incubated at room temperature with 50 μL of TruStain FcX (αCD16/CD32, 1:100, Biolegend #101319) for 20 minutes before staining with the Zombie UV Fixable Viability Stain (1:1000, Biolegend #423105). Samples were stained with the following for surface markers for 30 minutes at 4°C in different panels: CD45-PerCP (Clone 104), CD44-BV510 (clone IM7), Ly6C-BV570 (clone HK1.4), Ly6G-BV785 (clone 1A8), CD11b-BV650 (clone M1/70), CD11c-BV711 or BV421 (Clone N418), MHC-II I^A^/I^E^-FITC (clone M5/114.15.2), LAG3-PE (clone C9B7W), PD1-BV711 (clone 29F.1A12), CXCR3-PE/Fire810 (clone S18001A), CD107a-APC (clone AD4B), CD24-PE (clone M1/69), CD64-PE/Cy7 (clone X54-5/7.1), NK1.1-APC (clone PK136), CD103-APC/Cy7 (clone 2E7), (all Biolegend), CD45-BUV396 (Clone 104), CD4-BUV496 (clone GK1.5), CD8α-BUV73 (cloneH35-17.2), TCRβ-BUV615 (clone H57-597), CD25-FITC (clone 7D4), CD19-BUV737 (clone ID3), CD80-BUV563 (clone 16-10A1), CD86-BUV805 (clone PO3), F4/80-BUV395 (clone T45-2342) (all BD Biosciences), and Siglec F - NovaFluor Blue/610/70S (1RNM44N)(Invitrogen). For detection of intracellular Arginase-1, cells were then fixed and permeabilized using a Foxp3/Transcription Factor Staining buffer set (Thermo Fisher Scientific #00-5523-00) and then stained for Arginase 1-Alexa Fluor 700 (clone A1exF5; BD Biosciences). During analysis, debris, doublets, and dead cells were excluded before gating to identify monocytes (CD45^+^ CD11b^+^ Ly6C^+^ CD11c^-^ MHCII^-^), interstitial macrophages (IM; CD45^+^ CD11b^+^ Ly6C^+^ CD11c^-^ MHCII^+^), alveolar macrophages (AM, CD45^+^ CD11b^+^ Ly6C^+^ CD11c^+^ CD64^+^), and neutrophils (CD45^+^ CD11b^+^ Ly6G^+^). A complete gating strategy can be viewed in Supplemental Figure 1.

### Suppression assays

Spleens from naïve mice were harvested and processed into single-cell suspensions as described above. CD8a+ T cells were isolated via magnetic-activated cell separation (MACS, Miltenyi #130-104-075). Cells were stained with CellTrace Violet (Invitrogen #C34557) for 20 minutes at 37°C and resuspended in cRPMI supplemented with 5 ng/mL recombinant mouse IL-2 (Biolegend #575404). 5×10^5^ cells/well plated in a 96-well round-bottom plate. Mouse T-cell activator CD3/CD28 Dynabeads (Gibco # 11-452-D) were washed with PBS, resuspended in IL-2-supplemented cRPMI, and plated at a concentration of 5×10^5^ beads/well.

Lungs of influenza/A/X31-infected mice were harvested at 6 dpi and processed into homogenates. Suppressor cell populations were isolated via MACS technology using a mouse Myeloid-Derived Suppressor Cell Isolation Kit (Miltenyi 130-094-538), which purifies both Gr1^dim^Ly6G^-^ and Gr1^hi^Ly6G^+^ populations. Suppressor cells were resuspended in IL-2-supplemented cRPMI and plated in duplicate with the CD8 responder T cells at ratios of 1:1, 1:3, 1:9, 1:27, or 1:81.

An aliquot of each cell population was taken post-separation and stained for flow cytometry as described above to confirm purity of the isolation. Plates containing suppressor-responder co-cultures were incubated at 37°C with 5% CO_2_ for 48 hours. The cells were then centrifuged at 300 x g for 5 minutes and transferred to 96-well V-bottom plates before staining for flow cytometry using the method described above. To access the level of baseline activation and proliferation, wells containing unstimulated CD8 T cells only were used as a non-proliferation control, while wells containing stimulated CD8 T cells only were used as a proliferation control.

### Administration of Gemcitabine

Mice were i.p. injected with 12 mg/kg Gemcitabine (GEM, Sigma G6423) in a volume of 200 μL at 4 dpi. Mice were monitored for survival or euthanized at 6 dpi to confirm depletion efficacy by flow cytometry.

### Statistical analysis

For survival experiments, area under the curve (AUC) was analyzed by Mann-Witney U to test for differences in survival between different experimental groups. For flow cytometry comparisons, all data was analyzed via Mann-Whitney U test (for 2 groups) or Kruskal-Wallis test with Dunn’s Multiple Comparisons Test (for 3+ groups). A p-value of < 0.05 was considered statistically significant.

## Results

### Influenza/A/X31 infection protects against *Pb*E CD8-T cell induced MA-ARDS / MA-ALI

While mice infected with *Pb*E typically succumbed to infection beginning at day 6 post infection (dpi), mice co-infected with *Pb*E and influenza/A/X31 survived up to 27 dpi, ultimately succumbing to hyper-parasitemia (Figure 1A). This survival difference was not due to better control of parasite burden, as parasitemia was the same between the *Pb*E and co-infected groups. To test whether severe pulmonary vascular leak developed in the co-infected mice, we quantified lung permeability by three different methods (Figure 1B). Co-infected mice developed some vascular leak, as demonstrated by the capture of Evans blue dye from the lungs and the presence of protein in the BALF, however there was a trend towards co-infected mice having less vascular leak than the *Pb*E singly-infected mice. However, there was a significant protection against vascular leak in the co-infected group compared to the *Pb*E group with respect to the FITC-dextran permeability assay (Kruskall Wallis P<0.05)

**Figure 1.**
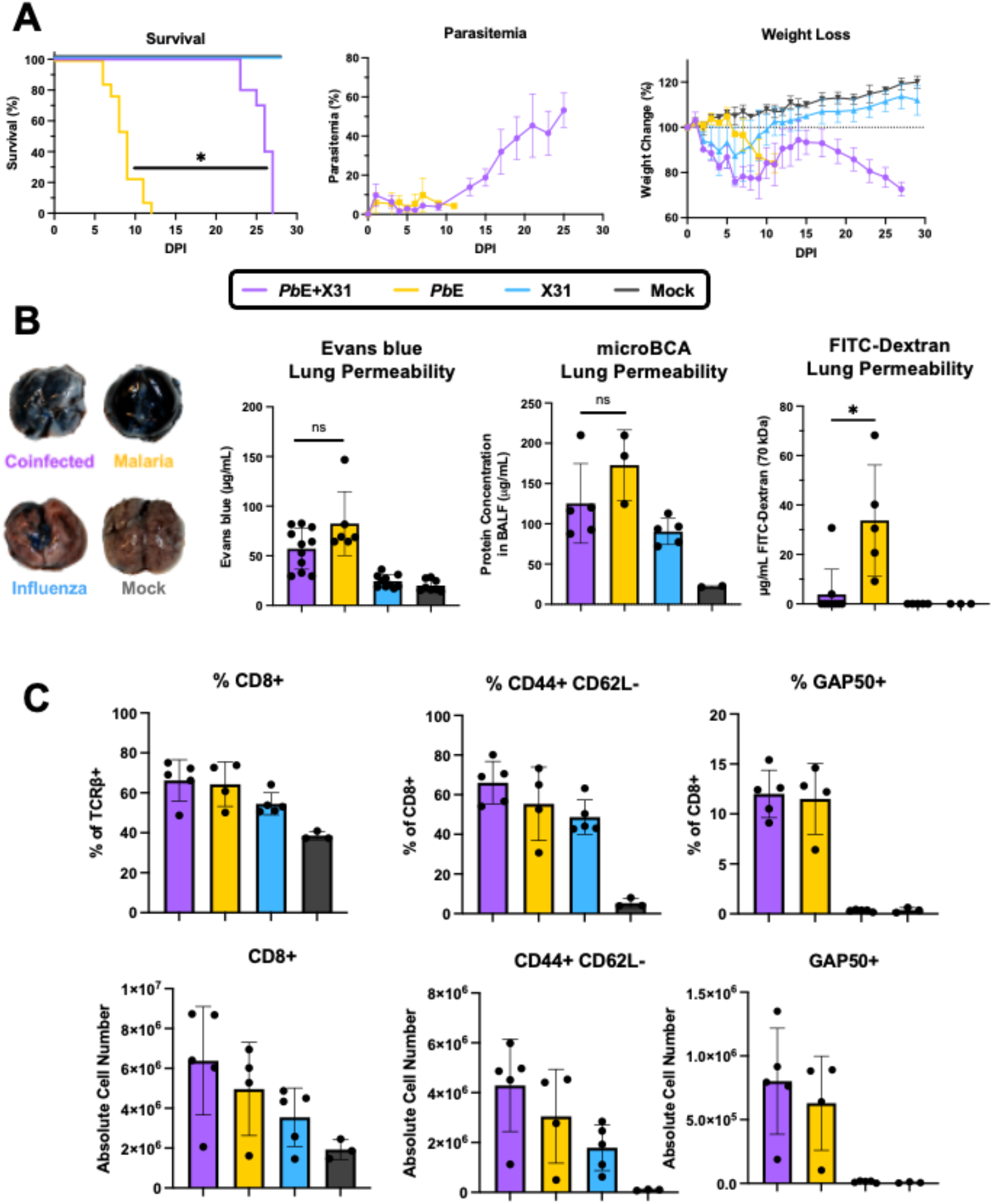
Co-infection with influenza/A/X31 prolongs survival during *Pb*E infection. (**A**) Mice co-infected with influenza/A/X31 and *Pb*E (purple line) survive significantly longer (Mann-Whitney U of AUC, p<0.0001) than those infected with *Pb*E only (yellow line), despite maintaining equivalent parasitemia and losing more weight. PbE+X31 n=10, PbE n=10, X31 n=10, Mock n=6. (**B**) Lung permeability was measured by Evans blue accumulation in the lungs (representative photos, left graph), protein concentration in the BALF as quantified by microBCA (middle graph), and FITC-dextran movement into the lungs from the blood (right graph). The co-infected and *Pb*E-only mice did not have statistically different levels of lung permeability by Evans blue (p>0.999) or microBCA (p>0.999), but the FITC-dextran assay suggested that they do not develop as severe lung permeability (p=0.013). Evans blue: PbE+X31 n=11, PbE n=6, X31 n=10, Mock n=8. microBCA: PbE+X31 n=5, PbE n=3, X31 n=5, Mock n=2. FITC-Dextran: PbE+X31 n=9, PbE n=5, X31 n=5, Mock n=3. (**C**) Co-infected and *Pb*E-only mice did not have significantly different frequencies nor absolute cell numbers of total CD8+ T cells (left, Mann-Whitney U, freq p>0.99, number p=0.29), CD44+CD62L-activated CD8 T cells (middle, Mann-Whitney U, freq p=0.69, number p=0.56), nor GAP50-reactive CD8 T cells (right, Mann-Whitney U, freq p=0.29, number p=0.56). * indicates P<0.05 by Kruskal Wallis with Dunn’s multiple comparison test. PbE+X31 n=5, PbE n=4, X31 n=5, Mock n=3.

Since vascular leak in *Plasmodium* infections is a CD8 T cell-driven immunopathology^20,21^ we used flow cytometry to examine the CD8 T cell response at 6 dpi, which is the critical window for MA-ALI/ MA-ARDS to begin developing. The frequency and absolute cell numbers of total and activated (CD44^+^ CD62L^-^) CD8 T cells responding to the lungs was similar between the co-infected and *Pb*E groups (Figure 1C). Additionally, the frequency of *Plasmodium*-glideosome associated protein 50 (GAP50)_40-48_^22^ -reactive CD8 T cells, as measured by tetramer staining, did not reveal any differences in the *Plasmodium*-reactive CD8 T cell response in the lung.

### Influenza/A/X31-induced protection only occurs during the innate phase of the immune response

Since parasite burden was not affected by coinfection, we hypothesized that the anti-influenza immune response may alter the kinetics of the pathogenic inflammatory anti-*Plasmodium* response in the lung. To determine if the innate or adaptive side of the anti-influenza immune response was involved in mediating influenza/A/X31-mediated survival of *Pb*E-infected mice, we altered the timing of the co-infection. Mice were infected with influenza/A/X31 either three days prior to (−3 dpi) or three days after (3 dpi) *Pb*E infection and monitored for survival. However, protection was only conferred when influenza/A/X31 infection was administered on the same day as PbE infection (Figure 2). The suggests that the anti-influenza innate immune response plays a significant role in protection, but that it also requires a certain timeframe in which to disrupt the developing immunopathology of MA-ARDS/MA-ALI.

**Figure 2.**
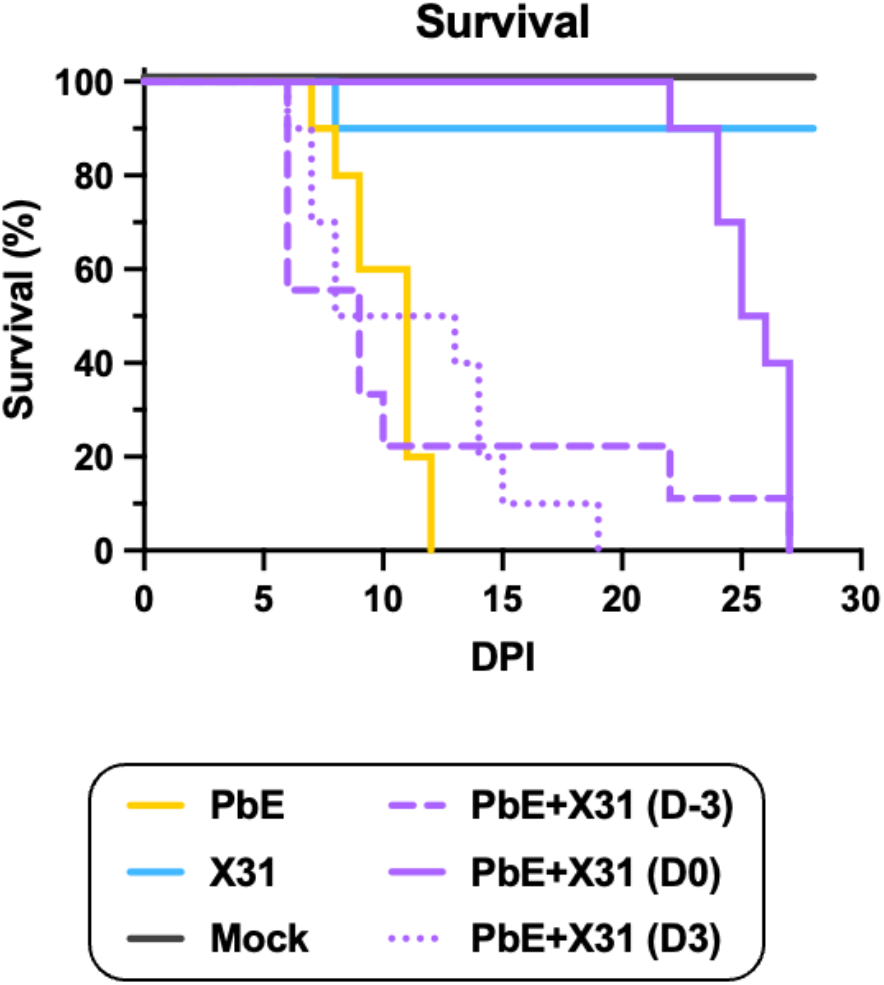
The timing of co-infection determines the impact on survival from MA-ALI / MA-ARDS. Influenza/A/X31 infections occurred at D-3 (dashed purple), D0 (solid purple), or D3 (dotted purple) compared to the *Pb*E infection. The D-3 and D3 co-infected groups succumbed to disease at a similar timeline to *Pb*E (yellow) (Kruskall-Wallis with Dunn’s Multiple Comparisons Test of AUC, D-3 vs *Pb*E p>0.999, D3 vs *Pb*E p>0.999), in contrast to the D0 co-infected group, which survived up to 27 dpi. PbE+X31 (D-3) n=9, PbE+X31 (D0) n=10, PbE+X31 (D3) n=10, PbE n=10, X31 n=10, Mock n=6.

### Influenza/A/X31 induced Arginase-1 expressing monocytes can suppress CD8 T cell responses

To determine which cells in the innate immune system were responsible for protection, we performed flow cytometry on the cells infiltrating the lung in response to influenza/A/X31-infection at 6 dpi. We quantified the phenotype of monocytes, IMs, AMs, and neutrophils that infiltrate the lung after influenza/A/X31-infection. Arginase 1 (Arg1) is a suppressive molecule expressed in both monocytes/macrophages and neutrophils that has been demonstrated to suppress CD8 T cell responses in tumors.^23,24^ We determined that influenza/A/X31 induces infiltration of Arg1-expressing AMs at 6 dpi (Figure 3A, 3B) but very few Arg1+ monocytes, IMs, and neutrophils present in the lung.

**Figure 3.**
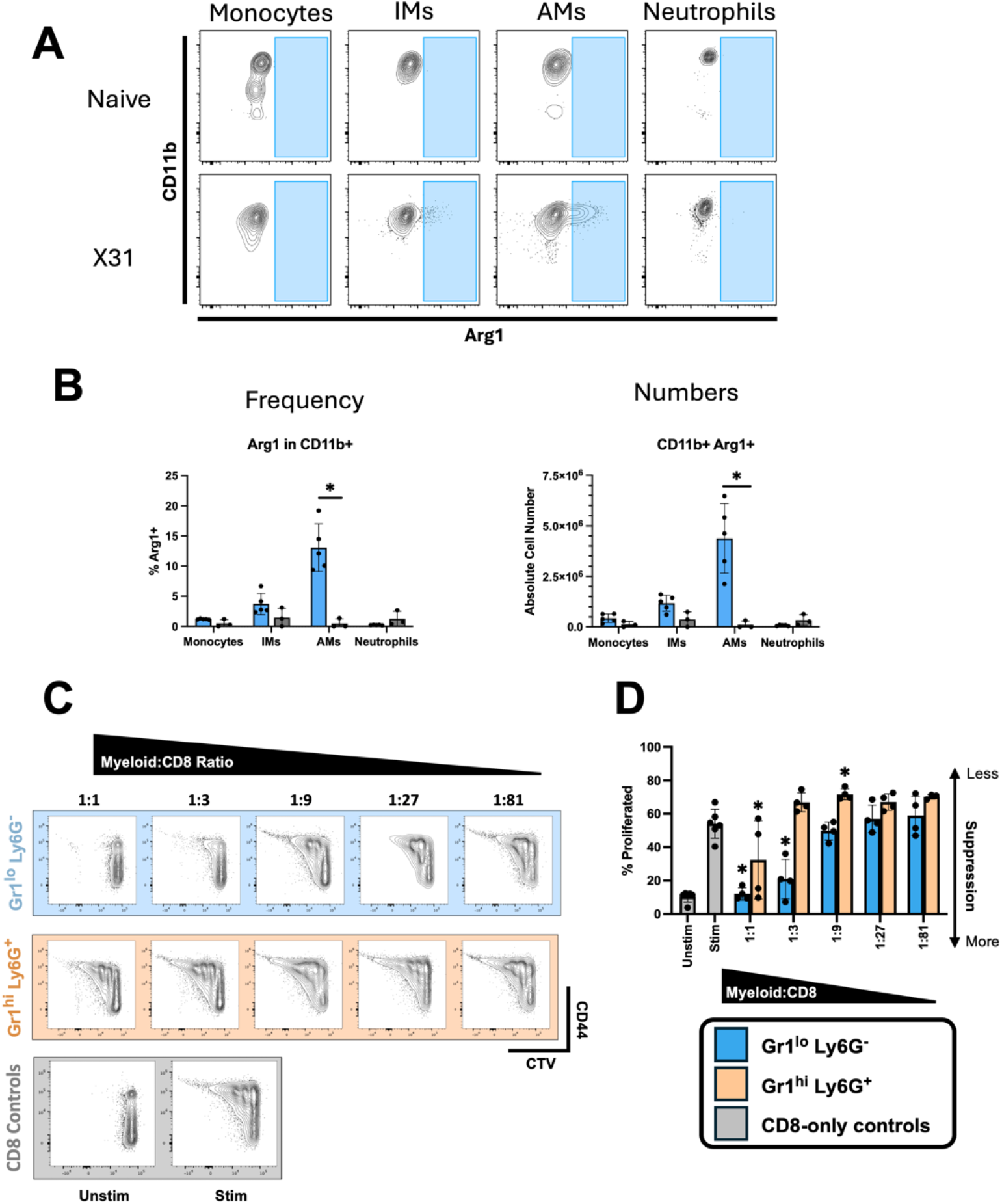
Influenza/A/X31 induces an Arg1+ AM response in the lungs which is capable of suppressing CD8 T cell activation and proliferation. (**A, B**) By flow cytometry at 6 dpi, we observed a significant increase in the frequency and number of lung Arg1+ AMs at 6 dpi during influenza/A/X31 infection compared to naïve (Mann-Whitney U, freq p=0.036, number p=0.036). In contrast, there was no increase in the frequency or number of Arg1+ monocytes (Mann-Whitney U, freq p=0.25, number p=0.071) Arg1+ IMs (Mann-Whitney U, freq p=0.25, number p=0.071) nor Arg1+ neutrophils (Mann-Whitney U, freq p=0.095, number p=0.095). * indicates P<0.05 by Mann-Whitney U test. X31 n=5, Mock n=3 (**C, D**) Gr1^lo^ Ly6G^-^ cells were able to suppress CD8 T cell proliferation *in vitro* at higher ratios (1:1, 1:3) and were statistically different from the CD8-only stimulated control group (mixed effects model, 1:1 p<0.0001, 1:3 p<0.001). These cells were better able to suppress than Gr1^hi^ Ly6G^+^ cells, which were only capable of suppression at the highest ratio (1:1 p=0.003) and promoted proliferation at the 1:9 ratio (p=0.01). * indicates a statistically significant difference (P<0.05) from the CD8-only stimulated group, by a mixed effects model. Gr1^lo^Ly6G^-^ n=4, Gr1^hi^Ly6G^+^ n=4, CD8-only n=6.

To demonstrate if Arg1+ AMs induced by influenza/A/X31 are capable of interfering with the pathogenic anti-*Plasmodium* CD8 T cell response in the lung, we performed an *in vitro* suppression assay. We isolated naïve CD8 T cells, stimulated them with anti-CD3 / CD28 beads and co-culturing them at different ratios of Gr1^dim^Ly6G^-^ (monocytes/macrophages) and Gr1^hi^Ly6G^+^ (neutrophils) isolated from the lungs of influenza/A/X31-infected mice at 6 dpi. As expected, CD8 T cells co-cultured with Gr1^hi^Ly6G^+^, which do not express Arg1 during influenza/A/X31 infection, were able to proliferate to the extent of CD8 T cells cultured with no myeloid cells as demonstrated by CTV staining and upregulated activation markers such as CD44 (Figure 3C, 3D). In contrast, the expansion and activation of CD8 T cells was inhibited when co-cultured with higher ratios (1:1 and 3:1) of Gr1^dim^Ly6G^-^ cells containing Arg1-expressing cells. This data provides a mechanism by which Arg1+ AM could suppress CD8 T cell-mediated MA-ALI / MA-ARDS in *Pb*E infection.

### Ly6C+ cells are required for protection against MA-ARDS/MA-ALI

To determine if the Arg1+ macrophages induced by influenza/A/X31 are required for the protective phenotype observed during co-infection, we depleted these cells using gemcitabine. This drug is a proposed cancer therapeutic that has been demonstrated to effectively deplete Ly6C^+^ cells.^25^ To measure the efficacy of the depletion, flow cytometry was used to analyze the immune cells present in the lungs at 6 dpi. While the Ly6C^+^ compartment was significantly decreased following gemcitabine treatment, the Ly6G^+^ compartment was unaffected (Figure 4A).

**Figure 4.**
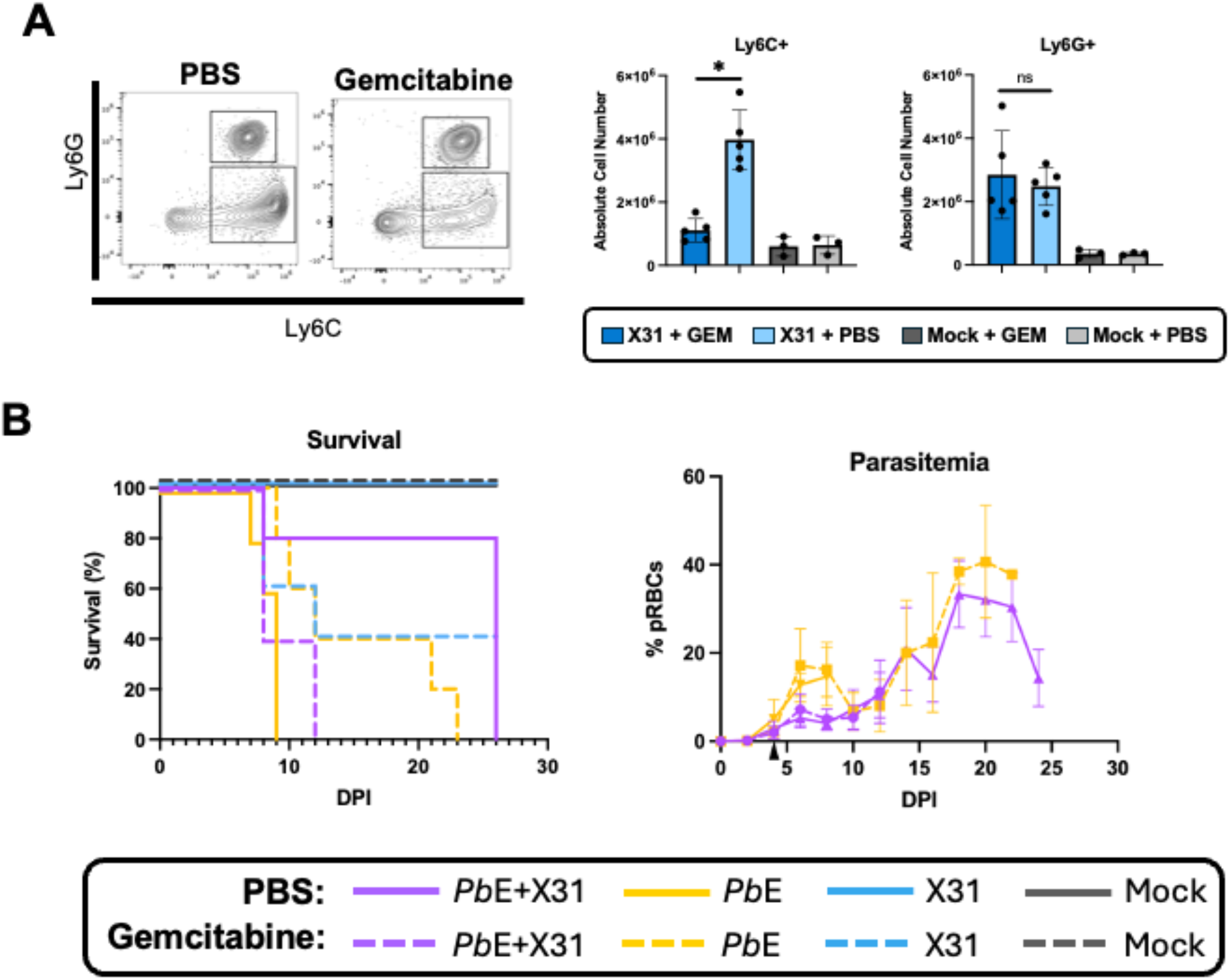
Gemcitabine treatment depletes Ly6C+ cells and reverts the coinfection survival phenotype without affecting parasitemia. (**A**) Mice treated with gemcitabine at 4 dpi had a decrease in the number of Ly6C+ cells responding to the lung compared to PBS-treated mice (Mann Whitney U, p=0.008) at 6 dpi. In contrast, the number of Ly6G+ cells was not affected by gemcitabine treatment (Mann Whitney U, p>0.999). * indicates a statistically significant (P<0.05) difference between the gemcitabine-treated vs. PBS-treated conditions. X31+GEM n=5, X31+PBS n=5, Mock+GEM n=3, Mock+PBS n=3. (**B**) Co-infected mice treated with gemcitabine succumbed to disease at a similar timeline to the *Pb*E-only mice (dashed purple vs. solid yellow). Gemcitabine treatment slightly prolonged death in the *Pb*E-only group (yellow dashed), while it resulted in a drop in survival from 100% to 40% in the influenza/A/X31-only group (blue, solid vs. dashed). Gemcitabine treatment did not exhibit any anti-parasitic effects, as exhibited by equivalent parasite burden in the gemcitabine-treated vs PBS-treated groups (right). PbE+X31+GEM n=5, PbE+X31+PBS n=5, PbE+GEM=5, PbE+PBS n=5, X31+GEM n=5, X31+PBS n=5, Mock+GEM n=3, Mock+PBS n=3.

We found that gemcitabine-treated co-infected mice died at the same rate as mice that had *Pb*E only, marking a complete reversion of the protective effects of Influenza A/X31 on survival (Figure 4B). We also observed no anti-parasitic effect of the drug, as parasite burden was the same between treated and untreated groups. Gemcitabine treatment slightly prolonged the survival of mice infected with *Pb*E only, suggesting that Ly6C^+^ cells may be mildly pathogenic in malaria. In contrast, the gemcitabine treatment caused the survival rate of mice singly-infected with influenza/A/X31 to drop from 100% to 40%. This observation reinforces the hypothesis that the Ly6C^+^ cells induced in the anti-influenza response are protective against severe disease, as the prognosis is worse when they are depleted.

## Discussion

This study identified a novel co-infection phenotype between *Pb*E and influenza/A/X31, wherein mice that are co-infected survive significantly longer than mice with malaria only (Figure 1). This phenotype results from differential immunopathology, rather than improving parasite control. We have identified a population of Arg1-expressing alveolar macrophages that respond to the lung following influenza/A/X31 coinfection (Figure 2) and are required for protection during co-infection (Figure 4). These cells are capable of suppressing CD8 T cell proliferation and activation (Figure 3), which may be relevant in disrupting MA-ARDS/MA-ALI, which are CD8 T cell-driven conditions. These cells are reminiscent of myeloid derived suppressor cells (M-MDSC) that have been well-characterized in tumor immunology and may represent a novel therapeutic target for organ-specific malaria pathologies in malaria.^26^

Previous work shows that alveolar macrophages both develop from fetal monocytes that differentiate into long-lived cells in the first week of life via GM-CSF^27^ and infiltrate from the bone marrow.^28^ In influenza infection, alveolar macrophages have been shown to be programmed by the gut microbiota influencing the severity of infection.^29^ On the other hand, alveolar macrophages have been shown to be depleted during *Plasmodium* infection and to repopulate by self-renewal.^30^ We hypothesize that Arg1+ alveolar macrophages we have quantified in influenza/A/X31 traffic from the bone marrow, as they are influenza-induced. This is similar to M-MDSCs in tumors which are thought to populate tumors via differentiation in the bone marrow and trafficking into the tumor environment.^31^

Additional research is needed to understand the mechanism underlying Arg1+ macrophage-driven protection against MA-ARDS/MA-ALI. While Arg1 levels are correlated with the level of protection, it remains unclear if the phenotype is dependent on Arg1 or if other mechanisms are involved. Suppressive macrophages in tumor environments have been reported to inhibit immune responses via numerous additional mechanisms, including the production of reactive oxygen species (ROS), inducible nitric oxide synthase (iNOS), and interleukin-10 (IL-10).^23–25^ By elucidating the mechanism through which these cells provide protection against CD8 T cell-induced vascular leak, other means of inducing those responses can be explored, opening the door for therapeutics to be developed that could work in conjunction with current anti-malarial treatment regimens.

It is surprising that CD8 T cell numbers are not altered by coinfection compared to *Pb*E alone (Figure 1), but the phenotype may be explained by endothelial activation. Upon activation, endothelial cells can secrete a variety of chemokines and cytokines that promote immune cell recruitment to the tissue. There is evidence that endothelial cells activate during MA-ARDS, as measured by elevated levels of endothelial products like vascular endothelial growth factor A (VEGF-A) and placental growth factor (PIGF), and this activation is dependent on CD8 T cells.^32^ We predict that the coinfected mice have increased lung inflammation due to the elevated immune cell response compared to *Pb*E alone, which would result in increased endothelial activation and allow for the recruitment of the protective Arg1+ cells. Monocytes have been reported to be able to modulate MA-ALI by clearing sequestered iRBCs away from the endothelial layer.^33^ As such, we predict that the protective phenotype observed during coinfection may in part be due to monocyte interactions with endothelial cells and iRBCs, however further investigation is needed to confirm this hypothesis.

## Acknowledgements

We thank the staff of the Office of Comparative Medicine for excellent animal husbandry.

## Author Contributions

JSR devised the experiments, performed the experiments, analyzed data, and wrote the paper; RN performed experiments and reviewed the paper; MD performed experiments and reviewed the paper; BDE and TJL devised the experiments, supervised the experiments, analyzed data, and wrote the paper.

## References

1. Van den Steen, P. E. et al. Pathogenesis of malaria-associated acute respiratory distress syndrome. Trends Parasitol. 29, 346–358 (2013).

2. Daneshvar, C. et al. Clinical and laboratory features of human Plasmodium knowlesi infection. Clin. Infect. Dis. Off. Publ. Infect. Dis. Soc. Am. 49, 852–860 (2009).

3. Alamer, E. et al. Dissemination of non-typhoidal Salmonella during Plasmodium chabaudi infection affects anti-malarial immunity. Parasitol. Res. 118, 2277–2285 (2019).

4. Blevins, L. K. et al. Coinfection with Streptococcus pneumoniae Negatively Modulates the Size and Composition of the Ongoing Influenza-Specific CD8+ T Cell Response. J. Immunol. 193, 5076–5087 (2014).

5. Edosomwan, E. U., Evbuomwan, I. O., Agbalalah, C., Dahunsi, S. O. & Abhulimhen-Iyoha, B. I. Malaria coinfection with Neglected Tropical Diseases (NTDs) in children at Internally Displaced Persons (IDP) camp in Benin City, Nigeria. Heliyon 6, e04604 (2020).

6. Hogan, B. et al. Malaria Coinfections in Febrile Pediatric Inpatients: A Hospital-Based Study From Ghana. Clin. Infect. Dis. Off. Publ. Infect. Dis. Soc. Am. 66, 1838–1845 (2018).

7. Sandlund, J. et al. Bacterial Coinfections in Travelers with Malaria: Rationale for Antibiotic Therapy. J. Clin. Microbiol. 51, 15–21 (2013).

8. Thompson, M. G. et al. Influenza and Malaria Coinfection Among Young Children in Western Kenya, 2009–2011. J. Infect. Dis. 206, 1674–1684 (2012).

9. Villarino, N. F. et al. Composition of the gut microbiota modulates the severity of malaria. Proc. Natl. Acad. Sci. 113, 2235–2240 (2016).

10. Daneshvar, C., William, T. & Davis, T. M. E. Clinical features and management of Plasmodium knowlesi infections in humans. Parasitology 145, 18–31 (2018).

11. van der Heyde, H. C., Gramaglia, I., Sun, G. & Woods, C. Platelet depletion by anti-CD41 (αIIb) mAb injection early but not late in the course of disease protects against Plasmodium berghei pathogenesis by altering the levels of pathogenic cytokines. Blood 105, 1956–1963 (2005).

12. Srivastava, K. et al. Platelet factor 4 mediates inflammation in experimental cerebral malaria. Cell Host Microbe 4, 179–187 (2008).

13. Nguee, S. Y. T., Júnior, J. W. B. D., Epiphanio, S., Rénia, L. & Claser, C. Experimental Models to Study the Pathogenesis of Malaria-Associated Acute Respiratory Distress Syndrome. Front. Cell. Infect. Microbiol. 12, 899581 (2022).

14. Chang, W. L. et al. CD8(+)-T-cell depletion ameliorates circulatory shock in Plasmodium berghei-infected mice. Infect. Immun. 69, 7341–7348 (2001).

15. Van den Steen, P. E. et al. Immunopathology and dexamethasone therapy in a new model for malaria-associated acute respiratory distress syndrome. Am. J. Respir. Crit. Care Med. 181, 957–968 (2010).

16. English, M. et al. Interobserver variation in respiratory signs of severe malaria. Arch. Dis. Child. 72, 334–336 (1995).

17. Anidi, I. U. et al. CD36 and Fyn Kinase Mediate Malaria-Induced Lung Endothelial Barrier Dysfunction in Mice Infected with Plasmodium berghei. PLoS ONE 8, e71010 (2013).

18. Shah, S. S., Fidock, D. A. & Prince, A. S. Hemozoin Promotes Lung Inflammation via Host Epithelial Activation. mBio 12, e02399–20 (2021).

19. Vandermosten, L. et al. Adrenal hormones mediate disease tolerance in malaria. Nat. Commun. 9, 4525 (2018).

20. Claser, C. et al. Lung endothelial cell antigen cross-presentation to CD8+T cells drives malaria-associated lung injury. Scopus OA2019 (2019).

21. Claser, C. et al. CD8+ T cells and IFN-γ mediate the time-dependent accumulation of infected red blood cells in deep organs during experimental cerebral malaria. PloS One 6, e18720 (2011).

22. Howland, S. W. et al. Brain microvessel cross-presentation is a hallmark of experimental cerebral malaria. EMBO Mol. Med. 5, 916–931 (2013).

23. Menjivar, R. E. et al. Arginase 1 is a key driver of immune suppression in pancreatic cancer. eLife 12, e80721 (2023).

24. Sosnowska, A. et al. Inhibition of arginase modulates T-cell response in the tumor microenvironment of lung carcinoma. OncoImmunology 10, 1956143 (2021).

25. Buchholz, S. M. et al. Depletion of Macrophages Improves Therapeutic Response to Gemcitabine in Murine Pancreas Cancer. Cancers 12, 1978 (2020).

26. Ostrand-Rosenberg, S., Lamb, T. J. & Pawelec, G. Here, There, and Everywhere: Myeloid-Derived Suppressor Cells in Immunology. J. Immunol. Baltim. Md 1950 210, 1183–1197 (2023).

27. Guilliams, M. et al. Alveolar macrophages develop from fetal monocytes that differentiate into long-lived cells in the first week of life via GM-CSF. J. Exp. Med. 210, 1977–1992 (2013).

28. Malainou, C., Abdin, S. M., Lachmann, N., Matt, U. & Herold, S. Alveolar macrophages in tissue homeostasis, inflammation, and infection: evolving concepts of therapeutic targeting. J. Clin. Invest. 133, (2023).

29. Ngo, V. L. et al. Intestinal microbiota programming of alveolar macrophages influences severity of respiratory viral infection. Cell Host Microbe 32, 335-348.e8 (2024).

30. Lai, S. M. et al. Organ-Specific Fate, Recruitment, and Refilling Dynamics of Tissue-Resident Macrophages during Blood-Stage Malaria. Cell Rep. 25, 3099-3109.e3 (2018).

31. Porembka, M. R. et al. Pancreatic adenocarcinoma induces bone marrow mobilization of myeloid-derived suppressor cells which promote primary tumor growth. Cancer Immunol. Immunother. CII 61, 1373–1385 (2012).

32. Pham, T.-T. et al. Pathogenic CD8+ T Cells Cause Increased Levels of VEGF-A in Experimental Malaria-Associated Acute Respiratory Distress Syndrome, but Therapeutic VEGFR Inhibition Is Not Effective. Front. Cell. Infect. Microbiol. 7, 416 (2017).

33. Lagassé, H. A. D. et al. Recruited monocytes modulate malaria-induced lung injury through CD36-mediated clearance of sequestered infected erythrocytes. J. Leukoc. Biol. 99, 659–671 (2016).

34. Pollenus, E. et al. CCR2 Is Dispensable for Disease Resolution but Required for the Restoration of Leukocyte Homeostasis Upon Experimental Malaria-Associated Acute Respiratory Distress Syndrome. Front. Immunol. 11, (2021).

