## Supplemental Figure 1 for "Influenza-induced alveolar macrophages protect against death by malaria-associated acute lung injury"

**Affiliations:**

### Supplemental Figure 1

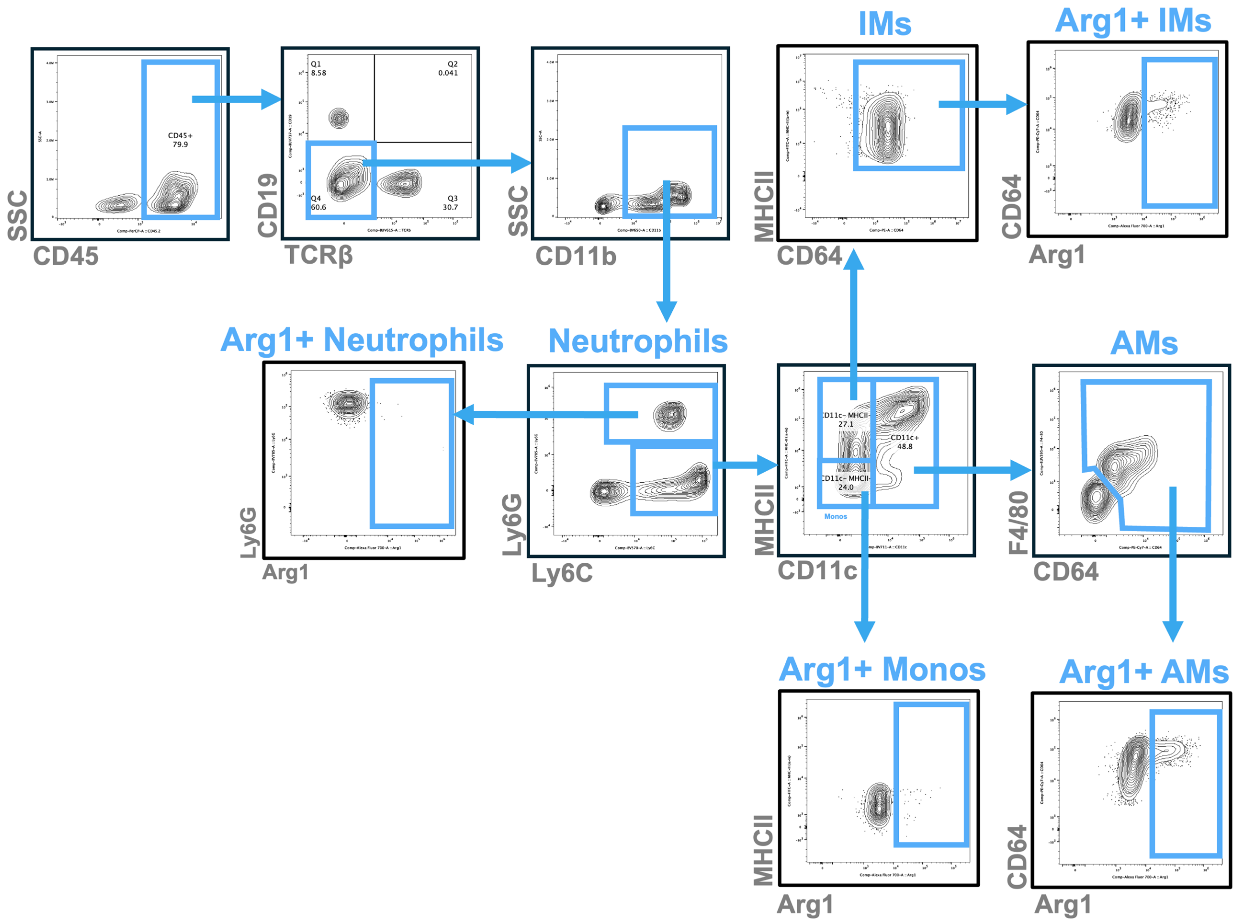


**Supplemental Figure 1. Gating strategy to identify lung myeloid cell subsets.** Debris, doublets, and dead cells were excluded (not shown) before selecting for CD45+ cells. B cells and T cells were excluded via CD19 and TCRβ. After gating on CD11b+ cells, Ly6C and Ly6G were compared to identify neutrophils (Ly6G+). From the Ly6C+ subset, CD11c and MHCII were compared. Monocytes were identified as CD11c- MHCII-. From the CD11c- MHCII+ population, IMs were identified as also being CD64+. From the CD11c+ population, AMs were identified as CD64+ F4/80+. Within each subset, Arg1 was examined to identify potential suppressor cells.
